# Screening of Stereochemically Defined 2,5-Diketopiperazines Identifies Autophagy Inducers without mTORC1 Suppression

**DOI:** 10.64898/2026.08.12.744315

**Authors:** Satoshi Yano, Shotaro Uchida, Shota Karakama, Shin Suzuki, Kuniki Kino, Taichi Hara

## Abstract

Modulating autophagy has emerged as a potential strategy for treating age-related diseases. However, commonly used pharmacological approaches to induce autophagy, particularly inhibition of mechanistic target of rapamycin complex 1 (mTORC1), can be associated with adverse effects, including immunosuppression and insulin resistance. This has prompted interest in autophagy modulators that act without directly inhibiting mTORC1. 2,5-Diketopiperazines (DKPs) are bioactive cyclic dipeptide scaffolds with diverse biological activities. However, systematic evaluation of their structure-activity relationships has been hindered by racemization during conventional chemical synthesis, leaving the contribution of stereochemistry to autophagy regulation poorly understood. Here, we used a stereoselective one-pot chemoenzymatic synthesis based on the adenylation domain of tyrocidine synthetase A to generate a DKP library with defined stereochemistry. Phenotypic screening in Caco-2 cells stably expressing the GFP-LC3-RFP autophagic flux probe identified four DKPs that increased autophagic flux: c(DW-DP), c(DW-LP), c(DF-DP), and c(DM-LP). Structure-activity analysis revealed stereochemistry-dependent effects associated with amino acid side-chain properties: D-configured residues were favored among DKPs containing aromatic amino acids or methionine, whereas L-configured residues were favored among those containing branched-chain amino acids. Substitution of the proline residue further altered activity, with glycine substitution tending to increase autophagic flux in some DKP scaffolds. Importantly, the active DKPs did not detectably reduce the phosphorylation of the mTORC1 downstream targets p70 S6K and 4EBP1, indicating that their autophagy-inducing effects do not require detectable suppression of canonical mTORC1 signaling. These findings establish stereochemically defined DKPs as candidate scaffolds for the development of autophagy inducers that act through mechanisms distinct from direct mTORC1 inhibition.

## 1. Introduction

Autophagy is a highly conserved, lysosome-dependent degradative pathway that maintains cellular homeostasis by turning over cytoplasmic constituents, including damaged organelles [1]. Accumulating evidence indicates that autophagic activity declines with age, and this decline is now recognized as a hallmark of aging, closely associated with impaired proteostasis and mitochondrial dysfunction [2]. Moreover, dietary restriction has been linked to lifespan extension in multiple model organisms, and these beneficial effects often depend, at least in part, on intact autophagy [3]. In parallel with dietary restriction, certain food-derived bioactive compounds have been implicated in the regulation of autophagy and promotion of longevity- and healthspan-associated phenotypes [4].

At present, the inhibition of mechanistic target of rapamycin complex 1 (mTORC1), most notably by rapamycin, is a widely used strategy for inducing autophagy and is also under investigation for therapeutic applications in selected settings [5]. However, direct mTOR inhibition can exert immunomodulatory and often immunosuppressive effects, as mTOR signaling regulates not only nutrient sensing but also T-cell activation and differentiation [6]. In addition, chronic mTOR inhibition may impair mTORC2 signaling, thereby contributing to insulin resistance, and has been associated with dyslipidemia in certain clinical settings [7, 8]. Together, these limitations highlight the need for autophagy enhancers that act through mechanisms distinct from canonical mTORC1 inhibition [9, 10].

Cyclic dipeptides, or 2,5-diketopiperazines (DKPs), are widely distributed in nature and occur in a variety of foods and beverages, including coffee, tea, fermented foods, and miso, supporting their relevance to orally consumed compounds [11, 12]. DKPs have attracted attention as bioactive scaffolds because their compact cyclic structure provides a conformationally constrained, stereochemically defined framework for side-chain presentation, making them attractive cores in medicinal chemistry [13, 14]. Consistent with this structural diversity, DKPs exhibit a wide range of biological activities, depending on their amino acid composition and stereochemistry. For example, cyclo(L-His-L-Tyr) has been reported to exhibit anticoagulant, chronotropic, and antibacterial activities, whereas DKPs containing D-amino acids differ markedly from their all-L counterparts in antimicrobial potency and activity profile [15-17].

In the context of autophagy-related biology, epidithiodiketopiperazine (ETP) alkaloids have been reported to induce autophagy in anticancer settings, suggesting that certain sulfur-containing DKP-related natural products can engage intracellular degradation pathways [18]. However, structure–activity relationship studies focusing on DKP-mediated modulation of autophagy remain limited. A major obstacle is synthetic accessibility, as conventional chemical routes often involve racemization during multistep synthesis, thereby hindering the large-scale preparation of stereochemically defined libraries. To address this challenge, we employed a one-pot chemoenzymatic synthesis method based on the adenylation domain of tyrocidine synthetase A (TycA-A), which was previously established by our collaborators [19]. By exploiting the stereorecognition properties of the enzyme, this method enables efficient DKP synthesis under mild aqueous conditions while minimizing racemization, thereby facilitating the preparation of structurally diverse DKPs with defined stereochemistry from various amino acid building blocks [19].

In this study, we constructed a stereochemically defined DKP library with systematically varied side-chain properties and screened it for autophagy-inducing activity using a GFP-LC3-RFP probe [20]. Cell imaging and quantitative analyses identified structural features associated with the activity, including side-chain composition, stereoselectivity, and effects of proline substitution. We also examined whether the active compounds acted independently of mTORC1 signaling. Collectively, these findings provide a basis for developing DKP-based autophagy inducers with a reduced risk of adverse effects.

## 2. Materials & Methods

### 2.1. Chemoenzymatic Synthesis of 2,5-Diketopiperazines (DKPs)

A library of 28 diketopiperazines (DKPs; Supplementary Figure S1A) and their substituted derivatives (Supplementary Figure S1B) was chemoenzymatically synthesized in a one-pot reaction using the adenylation domain of tyrocidine synthetase A (TycA-A), as described previously [19]. Briefly, this method enables intramolecular cyclization at high titers without the need for protection and deprotection steps and without racemization of substrate amino acids.

### 2.2. Preparation of Samples and Reagents

The DKP library and its substituted derivatives were prepared as previously described. For cell treatment, the compounds were dissolved in ultrapure water to prepare stock solutions, which were then diluted in the culture medium to a final concentration of 1 mM. Torin 1 and bafilomycin A1 were purchased from Merck KGaA (Darmstadt, Germany; 475991) and Cayman Chemical (Ann Arbor, MI, USA; 14005). Torin 1 and bafilomycin A1 were dissolved in DMSO to prepare stock solutions. Torin 1 and bafilomycin A1 were used at final concentrations of 1 μM and 200 nM, respectively, for 24 h as positive and negative controls for autophagic activity evaluation.

### 2.3. Cell Culture

The human epithelial colorectal adenocarcinoma cell line, Caco-2, was obtained from the ATCC (Manassas, VA, USA). Cells were cultured in DMEM (Fujifilm Wako Pure Chemical Corporation, Osaka, Japan; 044-29765) supplemented with 10% fetal bovine serum (FBS; Thermo Fisher Scientific Inc.), 1% penicillin-streptomycin (PS; Fujifilm Wako Pure Chemical Corporation; 168-23191), and 1% MEM non-essential amino acids (Fujifilm Wako Pure Chemical Corporation; 139-15651) at 37°C in a humidified atmosphere containing 5% CO_2_.

### 2.4. Establishment of Caco-2 Cells Expressing GFP-LC3-RFP

Caco-2 cells stably expressing GFP-LC3-RFP were generated using a retroviral vector. Retrovirus particles were produced by co-transfecting HEK293FT cells with pMRX-IP-GFP-LC3-RFP (RIKEN BRC DNA BANK; RDB14601), pCG-gag-pol, and pCG-VSVG using FuGENE HD (Promega, Madison, WI, USA; E2311) for 72 h [21]. The culture supernatant containing the viral particles was then collected. Caco-2 cells were incubated with virus-containing medium in the presence of polybrene (8 μg/mL) for 48 h. Infected cells were subsequently selected using a medium containing 5 μg/mL puromycin.

### 2.5. Determination of Cellular Autophagic Flux

Cellular autophagic flux was determined as previously described. Caco-2 cells expressing GFP-LC3-RFP were seeded in 24-well plates at a density of 6.0 × 10^4^ cells/well and pre-cultured for 24 h. The cells were then treated with 1 mM DKPs, 1 μM Torin 1, and/or 200 nM bafilomycin A1 for 24 h. After treatment, the cells were washed with PBS and incubated with 0.25% trypsin for 5 min. The cells were then collected, and the fluorescence intensities were measured using a SA3800 Spectral Analyzer (Sony Biotechnology, Tokyo, Japan). GFP and RFP fluorescence intensities were quantified, and autophagic activity was evaluated by calculating the relative GFP/RFP ratio.

### 2.6. Fluorescence Microscopy

Following treatment, Caco-2 cells expressing GFP-LC3-RFP were fixed with 2% paraformaldehyde and mounted using SlowFade Diamond Antifade Mountant (Thermo Fisher Scientific Inc.; S36963). Cellular fluorescence was visualized using a confocal laser scanning microscope (FV3000; Olympus Corporation, Tokyo, Japan).

### 2.7. Western Blotting

Cells were lysed in Tris-Triton buffer containing 50 mM Tris-HCl (pH 7.4), 150 mM NaCl, 1 mM EDTA, 1% (v/v) Triton X-100, and a cOmplete protease inhibitor cocktail (Merck KGaA, Darmstadt, Germany; 04693116001). Cell lysates were centrifuged at 15,000 × *g* for 15 min at 4°C, and the supernatants were collected. Protein concentrations were determined using a Protein Assay Bicinchoninic Acid Assay Kit (Fujifilm Wako Pure Chemical Corporation). Equal amounts of protein were separated by SDS-PAGE and transferred onto mini 0.2 μm polyvinylidene difluoride transfer packs (Bio-Rad, Hercules, CA, USA; #1704156) using a Trans-Blot Turbo Transfer System (Bio-Rad). The membranes were blocked with 5% nonfat dry milk in TBST buffer (500 mM NaCl, 20 mM Tris-HCl, pH 7.4, and 0.1% Tween 20) and incubated overnight at 4°C with specific primary antibodies (1:1000 dilution), followed by horseradish peroxidase-conjugated secondary antibodies (1:10,000 dilution) for 1 h at room temperature. Bound antibodies were detected using the ECL system, and relative protein levels were quantified using a FUSION SOLO S (Vilber Lourmat, Marne-la-Vallée, France). The primary antibodies used were phospho-p70 S6K (#97596) and 4EBP1 (#9644), both purchased from Cell Signaling Technology, Inc. (Danvers, MA, USA). Pan-actin (C4) antibody (#41185; Cell Signaling Technology Inc., Danvers, MA, USA) was used as an internal control. Horseradish peroxidase-conjugated anti-mouse and anti-rabbit IgGs (115-035-003 and 111-035-144, respectively) were purchased from Jackson ImmunoResearch Laboratories (West Grove, PA, USA).

### 2.8. Statistical Analysis

Data are presented as the mean ± standard error of the mean (SEM). Statistical analyses were performed using a two-tailed Student’s *t*-test or one-way analysis of variance (ANOVA) for multiple comparisons. Differences were considered statistically significant at \**p* < 0.05, \*\**p* < 0.01, and \*\*\**p* < 0.001. Different letters above the bars indicate significant differences (*p* < 0.05). Heatmaps were generated using the R software.

## 3. Results

### 3.1. Screening of a Synthetic DKP Library Identifies Novel Autophagy Inducers

To identify novel autophagy-inducing compounds, we established a screening system using Caco-2 cells stably expressing the GFP-LC3-RFP probe. Prior to screening, the validity of the assay was quantitatively confirmed: treatment with Torin 1, an autophagy inducer, decreased the GFP/RFP ratio, whereas treatment with bafilomycin A1, an autophagy inhibitor, increased this ratio (Supplementary Figure S2C). Using this evaluation system, the autophagy-inducing activities of a library of 28 diketopiperazines (DKPs) synthesized via a chemoenzymatic method (Supplementary Figure S1A) were analyzed. This screening enabled a quantitative comparison of the effects of individual DKP compounds on autophagic flux.

Among the 28 DKPs in the library, four compounds were identified as autophagic flux enhancers. Quantitative analysis by flow cytometry revealed that the GFP/RFP ratio significantly decreased to below 0.9 in cells treated with c(DW-DP), c(DW-LP), c(DF-DP), and c(DM-LP) (Figure 1A). Furthermore, confocal laser scanning microscopy demonstrated that cells treated with these four compounds exhibited an increased number of puncta, reflecting the quenching of GFP fluorescence, similar to that observed in Torin 1-treated cells (Figure 1B and S2A; scale bar = 10 μm). These results confirmed that the four DKPs can induce autophagy in cells.

**Figure 1.**
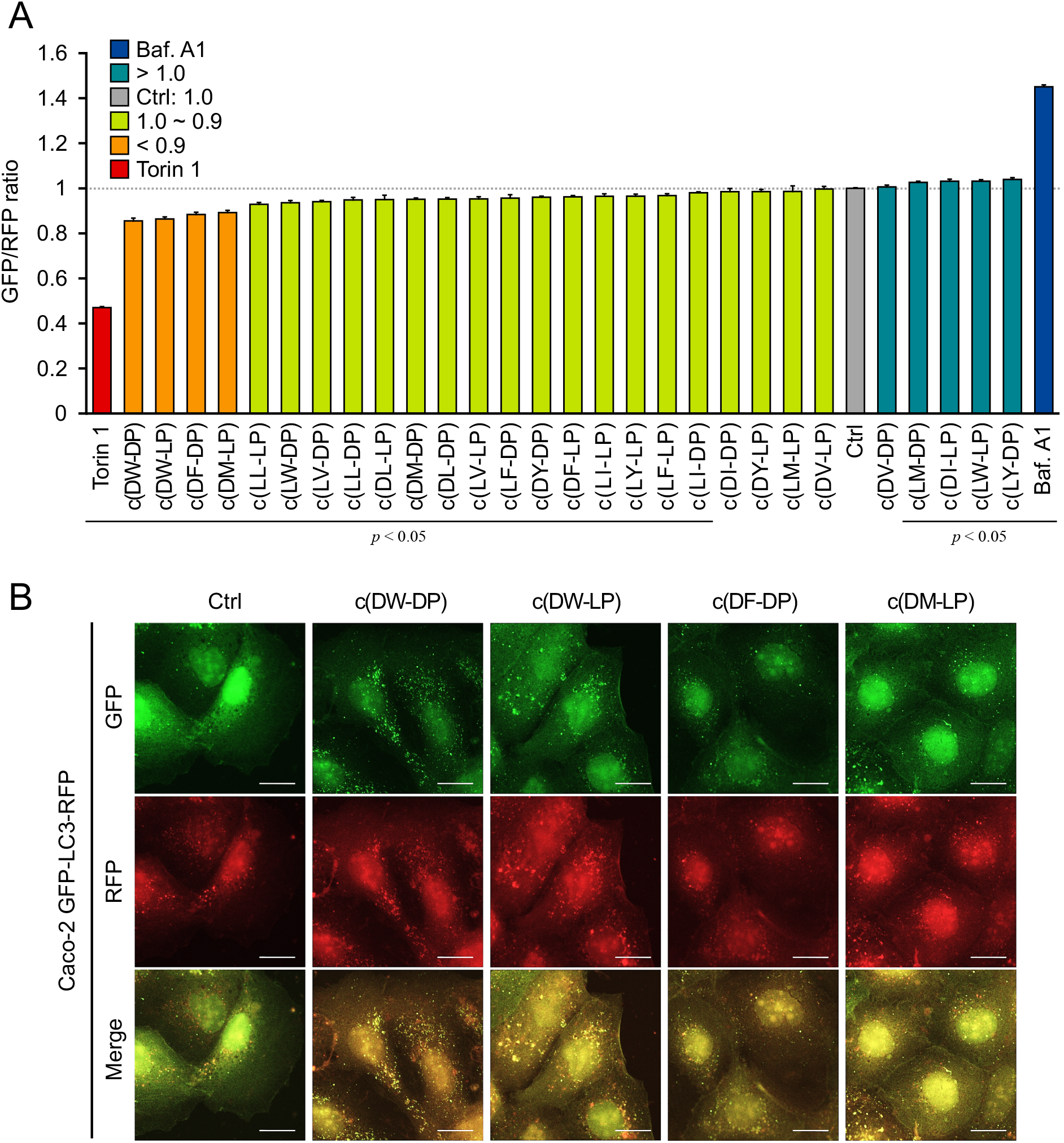
Screening of a synthetic DKP library identified autophagy-inducing compounds. (A) Quantitative analysis of autophagic flux in Caco-2 cells stably expressing GFP-LC3-RFP. Cells were treated for 24 h with 1 mM of each chemoenzymatically synthesized diketopiperazine (DKP), 1 μM Torin 1, or 200 nM bafilomycin A1 (Baf. A1), and the GFP/RFP ratio was measured by flow cytometry. Compounds that reduced the ratio to < 0.9 are highlighted in orange. (B) Representative confocal images of cells treated with vehicle control (Ctrl), c(DW-DP), c(DW-LP), c(DF-DP), or c(DM-LP). Increased puncta and reduced GFP fluorescence indicate autophagy. Scale bars, 10 μm. Data are presented as mean ± SEM (n=6). *p* < 0.05.

### 3.2. Autophagy-Inducing Activity of DKPs Depends on Specific Combinations of Amino Acid Side Chains and Stereochemistry

To elucidate the structural features of the identified active compounds, we analyzed the effects of the stereochemistry of the constituent amino acids on autophagy induction across the entire DKP library. The GFP/RFP ratios of the 28 compounds were visualized using heatmaps (Figure 2A). A comparison of the entire library according to the L- or D-form of the proline residue (Figure 2B) and according to the L- or D-form of the X-side amino acid (Figure 2C) revealed no statistically significant differences in the mean GFP/RFP ratios. These results indicate that the stereochemistry of a single amino acid does not account for the rules underlying the autophagy-inducing activity of DKPs.

**Figure 2.**
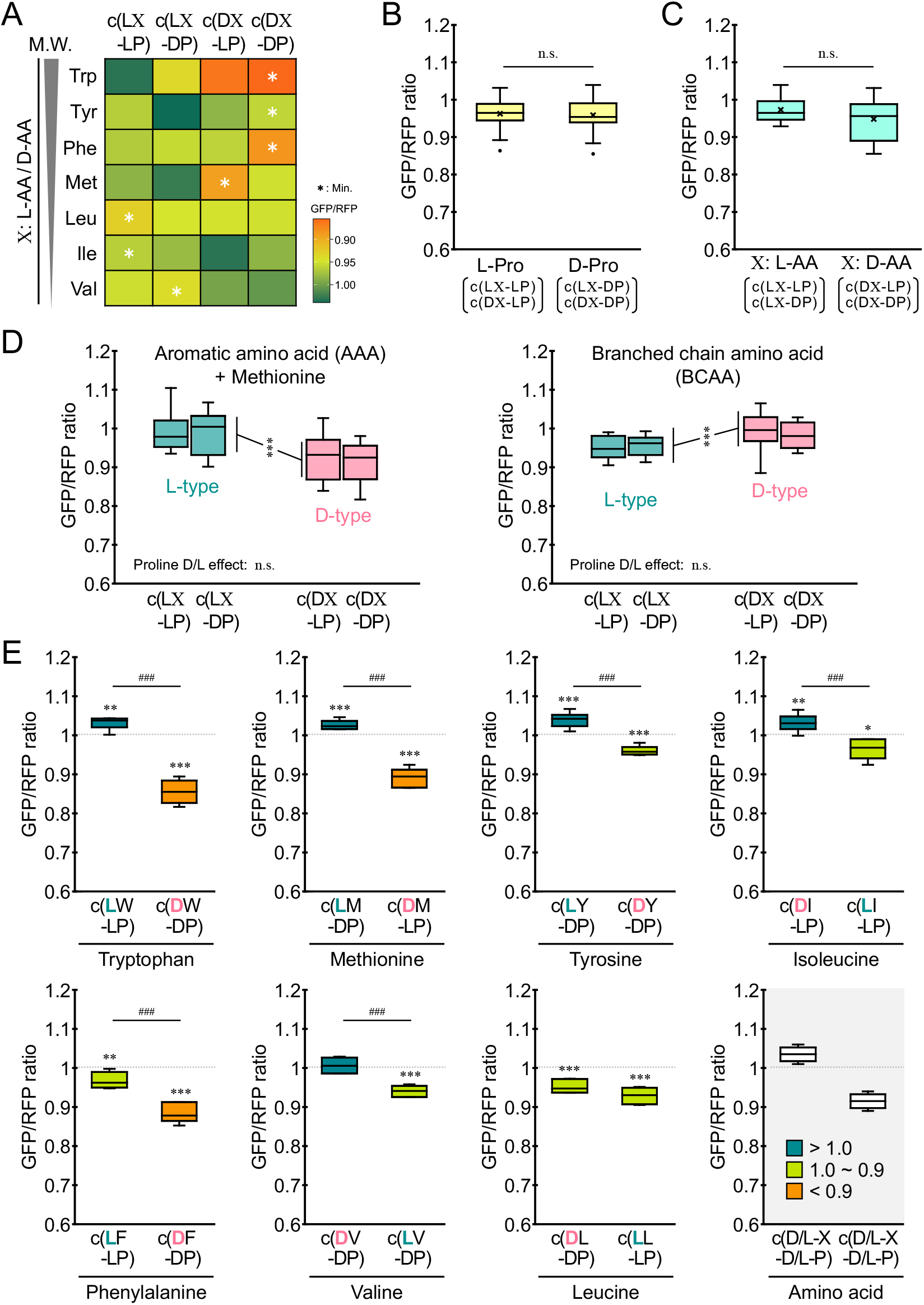
Autophagy-inducing activity of DKPs depends on amino acid side-chain properties and stereochemistry. (A) Heatmap of GFP/RFP ratios for 28 DKPs. Asterisks indicate the minimum GFP/RFP ratio in each amino acid group. (B, C) Mean GFP/RFP ratios for compounds grouped according to the stereochemistry of the proline residue (B) or the X-side amino acid (C). n.s., not significant. (D) GFP/RFP ratios stratified by the side-chain class of the X-side amino acid: aromatic amino acid and methionine (AAA + Met) group or branched-chain amino acid (BCAA) group. (E) Effects of stereochemistry on autophagy-inducing activity at the individual amino acid level (Trp, Met, Tyr, Ile, Phe, Val, and Leu). Data are presented as mean ± SEM (n=6). Statistical significance was determined using the two-tailed Student’s *t*-test or one-way analysis of variance (ANOVA). \**p* < 0.05, \*\**p* < 0.01, and \*\*\**p* < 0.001; ^###^*p* < 0.001, significant differences between specific structural pairs.

When the compounds were stratified according to the side-chain characteristics of the constituent amino acids, the specific stereoselectivity of the activity became apparent. The X-side amino acids were classified into an “aromatic amino acid and methionine (AAA + Met) group” and a “branched-chain amino acid (BCAA) group.” In the AAA + Met group, DKPs containing the D-form of the X-side amino acid significantly decreased the GFP/RFP ratio, whereas in the BCAA group, DKPs containing the L-form significantly decreased the ratio (Figure 2D). At the individual amino acid level, significant differences were observed, such that the D-form contributed to autophagy induction in Trp, Tyr, Phe, and Met, whereas the L-form contributed to Leu, Ile, and Val (Figure 2E). Together, these data indicate that the combination of amino acid side-chain properties and specific stereochemistry is a key determinant of autophagy induction.

### 3.3. Substitution of the Proline Residue Modulates the Autophagy-Inducing Activity of DKP Derivatives

To evaluate the structural contribution of the second residue forming the backbone of the active DKPs, we analyzed the autophagy-inducing activities of derivatives in which the proline residue was replaced with other amino acids (Supplementary Figure S1B). Based on the highly active compounds, the GFP/RFP ratios of the derivative groups in which the proline residue was substituted with Gly, trans-4-hydroxy-L-Pro (HYP), L-Ser, or D-Ser were evaluated. In some derivatives based on c(DW-LP) and c(DW-DP), and c(DF-DP), the reduction in the GFP/RFP ratio was attenuated (Figure 3A and 3B). The Gly-substituted derivative of c(DW-LP) and c(DW-DP) showed a tendency toward a greater reduction in the GFP/RFP ratio (*p* = 0.072, Figure 3A). In contrast, all substituted derivatives based on c(DM-LP) significantly enhanced the reduction in the GFP/RFP ratio compared to the original compound (Figure 3C). These findings suggest that substituting the second residue in DKPs with other amino acids can modulate their autophagy-inducing activity.

**Figure 3.**
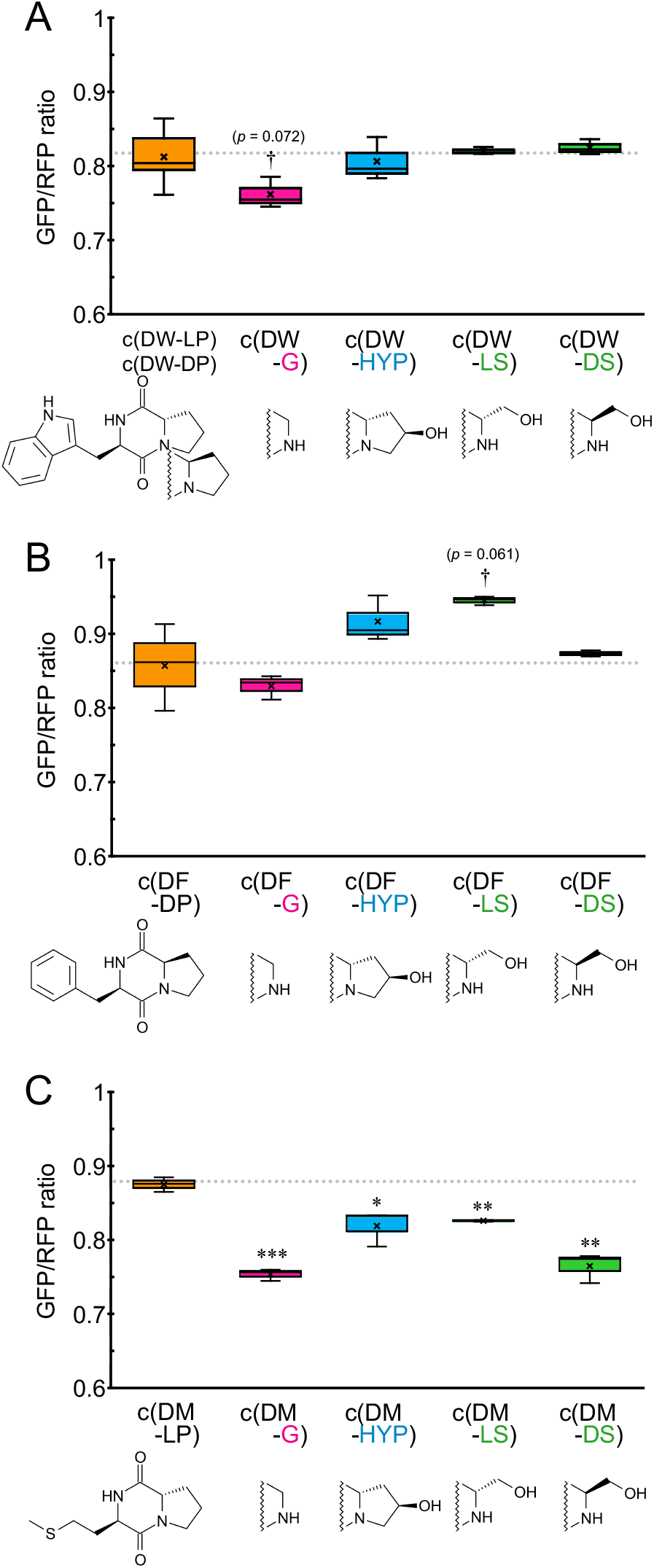
Substitution of the proline residue modulates the autophagy-inducing activity of DKP derivatives. (A–C) GFP/RFP ratios in Caco-2 cells treated for 24 h with the derivatives of active DKPs. The proline residue was replaced with Gly (-G), trans-4-hydroxy-L-Pro (-HYP), L-Ser (-LS), or D-Ser (-DS) residues. The derivatives of c(DW-DP) and c(DW-LP) are shown in panel A, those of c(DF-DP) in panel B, and those of c(DM-LP) in panel C. Data are presented as mean ± SEM (n=3). \**p* < 0.05, \*\**p* < 0.01, and \*\*\**p* < 0.001, versus the corresponding parent compound. *p* values of 0.072 and 0.061 indicated nonsignificant trends toward lower GFP/RFP ratios.

### 3.4. Active DKPs Induce Autophagy Independently of the mTORC1 Signaling Pathway and Exhibit Additive Effects with Torin 1

The effects of the identified active DKPs on the mTORC1 pathway, a major nutrient-sensing signaling pathway, were investigated. Using cells treated with the four active compounds, the phosphorylation levels of p70 S6K and 4EBP1, which are downstream targets of mTORC1, were analyzed using western blotting. While the Torin 1-treated group showed a marked decrease in the phosphorylation of both proteins, the DKP-treated groups showed no reduction in phosphorylation levels relative to the control group (Figure 4A). These results indicate that active DKPs induce autophagy without inhibiting mTORC1 signaling.

**Figure 4.**
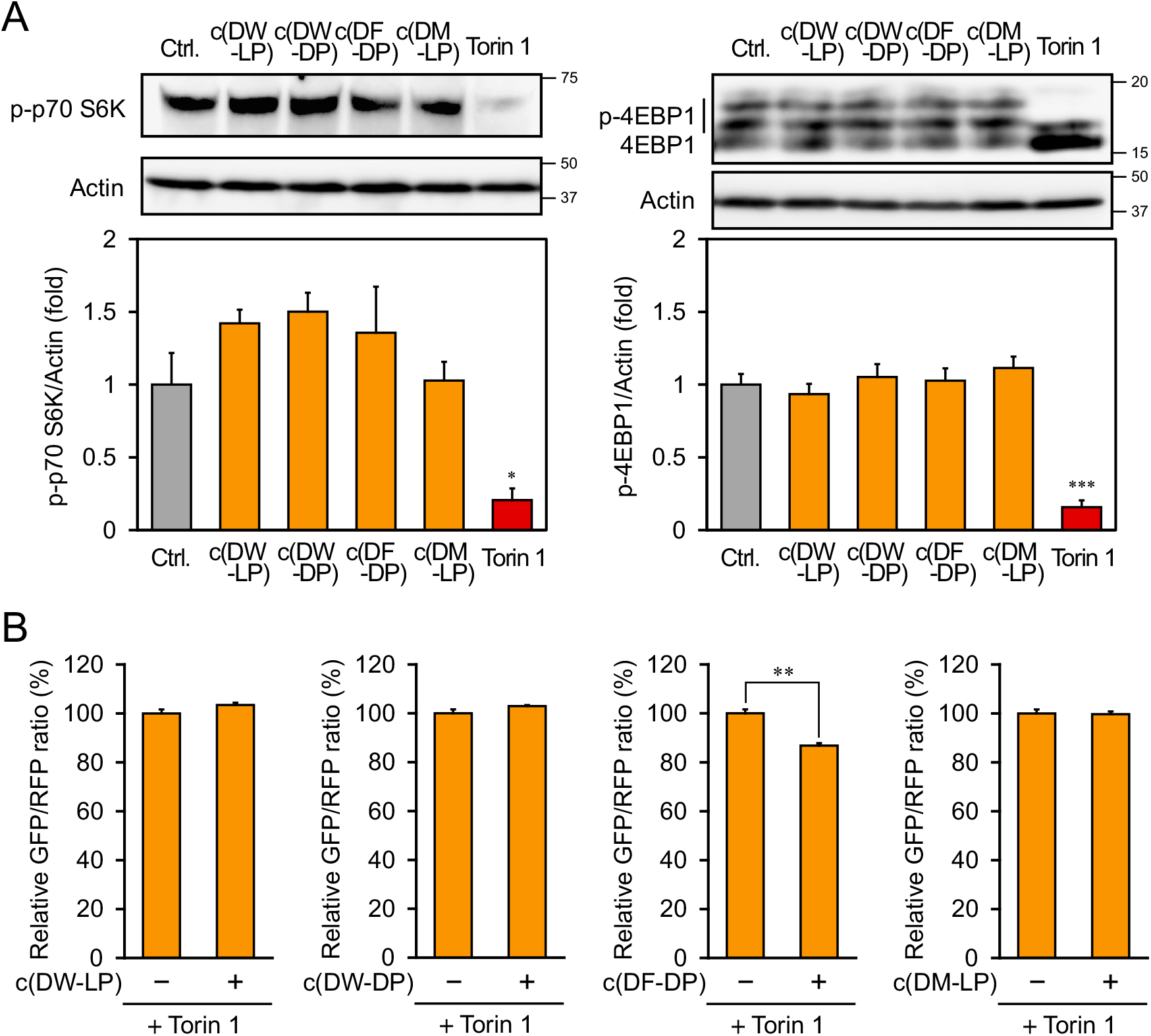
Active DKPs induce autophagy independently of mTORC1 inhibition and show additive effects with Torin 1. (A) Western blot analysis of mTORC1 downstream targets in Caco-2 cells treated for 24 h with 1 mM active DKPs or 1 μM Torin 1. Phosphorylated p70 S6K (p-p70 S6K) and 4EBP1 (p-4EBP1) were analyzed using pan-actin as a loading control. Bar graphs show phosphorylated protein levels normalized to actin. (B) GFP/RFP ratios measured by flow cytometry in cells co-treated for 24 h with each active DKP (1 mM) and Torin 1 (1 μM). Data are presented as mean ± SEM (n=3). \*\**p* < 0.01 versus Torin 1 alone.

Given that their mode of action differed from that of mTORC1 inhibition, the effect of co-treatment with DKPs and Torin 1 on autophagy induction was evaluated. The GFP/RFP ratio was measured in cells co-treated with each of the four active DKPs and Torin 1. The results showed that the GFP/RFP ratio was significantly reduced in the co-treatment group containing c(DF-DP) compared to the Torin 1-only treatment group (Figure 4B). This additive effect supports the notion that active DKPs promote autophagy through a pathway independent of canonical mTORC1 inhibition.

## 4. Discussion

In the present study, autophagy screening identified four DKPs—c(DW-DP), c(DW-LP), c(DF-DP), and c(DM-LP)—that induced autophagy without detectable suppression of canonical mTORC1 signaling. Analysis of autophagic activity revealed clear stereoselective patterns: the D-form was favored in DKPs containing aromatic amino acids or methionine, whereas the L-form was favored in those containing branched-chain amino acids. These findings support the idea that DKPs act through a mechanism distinct from direct mTORC1 inhibition, although the upstream target(s) remain to be identified.

Beyond their newly identified role in autophagy induction, the four active DKPs identified in this study have been associated with diverse physiological and pharmacological effects, further highlighting the broad functional potential of the DKP scaffold. For example, c(DF-DP) has been reported to exhibit antibacterial activity against Vibrio anguillarum [16]. In contrast, the all-L isomer cyclo(L-Phe-L-Pro) has been reported to exhibit cytotoxicity toward selected human tumor cell lines [22] and functions as a quorum-sensing signal in Vibrio spp. [23]. Similarly, tryptophan-containing DKPs, including c(DW-DP) and c(DW-LP), belong to cyclo(Trp-Pro) stereoisomers that modulate cardiac electrophysiology and heart rate in a stereochemistry-dependent manner [24]. These isomers have also been associated with lipid peroxidation and other stress-related changes in hepatocyte-based assays, with isomer-dependent differences observed in vivo [25]. Although the D-Met-L-Pro stereoisomer remains poorly characterized, related cyclo(Met-Pro) frameworks occur in probiotic- and food-associated cyclic dipeptides, supporting their relevance to orally consumed biological materials [26]. Together, these observations suggest that DKP stereoisomers engage diverse biological targets. Our finding that these compounds also induce autophagy without clear evidence of canonical mTORC1 suppression identifies an additional biological property of this scaffold and further supports the need for careful evaluation of their in vivo safety.

The dependence of activity on both amino acid side chain properties and stereochemistry suggests that DKPs are recognized stereoselectively by intracellular target proteins. Consistent with this idea, studies of natural DKP derivatives have shown that D-amino-acid-containing diastereomers and enantiomers can markedly alter minimum inhibitory concentrations and activity profiles relative to their all-L counterparts, indicating strict stereodiscrimination by biological macromolecules [16, 17]. In addition to amino acid chirality, the three-dimensional structure of DKPs can be influenced by the conformational constraints imposed by proline residues. Proline is known to restrict backbone conformational flexibility, and such conformational preorganization can influence molecular recognition and binding energetics in proline-containing peptides [27, 28]. However, whether this principle also applies to cyclic DKPs remains unclear. In the present study, replacement of the proline residue attenuated the activity of some derivatives, whereas glycine substitution and several c(DM-LP)-based derivatives tended to enhance activity, although these differences were not statistically significant. Thus, rather than indicating a general requirement for proline, our findings suggest that the conformational and steric properties imparted by the second residue may modulate the autophagy-inducing activity of DKPs in a scaffold-dependent manner. Further structural and target-binding studies will be required to determine whether proline-mediated conformational preorganization directly contributes to molecular recognition by active DKPs.

The finding that the identified DKPs enhanced autophagy without detectable suppression of mTORC1 signaling suggests that their mechanism of action differs from that of direct mTOR inhibitors such as Torin 1. Chronic pharmacological inhibition of mTOR signaling can produce undesirable effects, including impaired immune function through suppression of T-cell proliferation [6] and insulin resistance associated with unintended inhibition of mTORC2 [7]. Therefore, pharmacological induction of autophagy without direct mTORC1 inhibition may provide an alternative strategy for avoiding some of the liabilities associated with conventional mTOR-targeting approaches. Indeed, small-molecule autophagy enhancers acting independently of or downstream of mTOR have shown beneficial effects in experimental disease models [29], and the development of more selective autophagy modulators has been highlighted as an important direction for autophagy-targeted therapy [30]. Our stereoselective synthetic platform, which enables systematic functional screening of structurally defined DKP libraries, may facilitate the identification and optimization of such modulators.

Several limitations of this study should be acknowledged. Although the additive effect observed for c(DF-DP) in combination with Torin 1 is consistent with a mechanism distinct from direct mTOR inhibition, it does not establish pathway independence. Further mechanistic studies, including genetic or pharmacological epistasis-based approaches, will therefore be required. In addition, because the present screening was phenotype-based, the direct intracellular targets of the active DKPs remain unknown and will need to be identified through target-deconvolution approaches [31]. Finally, the in vivo efficacy, pharmacokinetic properties, and safety of the active DKPs remain to be evaluated in appropriate animal models.

Future studies should focus on the rational optimization of DKP derivatives based on the structural features identified in this study, along with the identification of their direct molecular targets. One potential candidate is the aryl hydrocarbon receptor (AhR). D-amino acids, including D-tryptophan, can be converted into potent AhR agonists through oxidative deamination by endogenous enzymes, followed by spontaneous condensation [32]. Appropriate activation of AhR signaling can regulate cellular stress responses and metabolic programs and may intersect with autophagy in a context-dependent manner. In summary, DKPs with defined side-chain properties and stereochemistry provide a promising molecular basis for developing mTORC1-independent autophagy inducers with improved safety profiles.

## Abbreviations

AAA: aromatic amino acids
AhR: aryl hydrocarbon receptor
ANOVA: analysis of variance
Baf. A1: bafilomycin A1
BCAA: branched-chain amino acids
DKP: 2,5-diketopiperazine
DMEM: Dulbecco’s modified Eagle’s medium
DMSO: dimethyl sulfoxide
ECL: enhanced chemiluminescence
EDTA: ethylenediaminetetraacetic acid
ETP: epidithiodiketopiperazine
FBS: fetal bovine serum
GFP: green fluorescent protein
HYP: trans-4-hydroxy-L-proline
IgG: immunoglobulin G
LC3: microtubule-associated protein 1 light chain 3
MEM: minimum essential medium
mTOR: mechanistic target of rapamycin
mTORC1: mechanistic target of rapamycin complex 1
mTORC2: mechanistic target of rapamycin complex 2
n.s.: not significant
PBS: phosphate-buffered saline
PS: penicillin-streptomycin
RFP: red fluorescent protein
SDS-PAGE: sodium dodecyl sulfate-polyacrylamide gel electrophoresis
SEM: standard error of the mean
TBST: Tris-buffered saline containing Tween 20
TycA-A: adenylation domain of tyrocidine synthetase A
4EBP1: eukaryotic translation initiation factor 4E-binding protein 1
p70 S6K: p70 ribosomal S6 kinase

## Acknowledgments

The authors thank the members of the Hara Lab and the Kino Lab for helpful suggestions and discussions.

## Author contributions

Conceptualization: S.Y., K.K., T.H.; Data curation and formal analysis: S.Y., S.S.; Investigation: S.Y., S.U., S.K.; Writing original draft: S.Y.; Supervision: K.K., T.H; Writing – review and editing: S.Y., K.K., T.H.

## Data availability

All data that support the findings of this study are available from the corresponding author upon reasonable request.

---

**Supplementary Figure S1.**
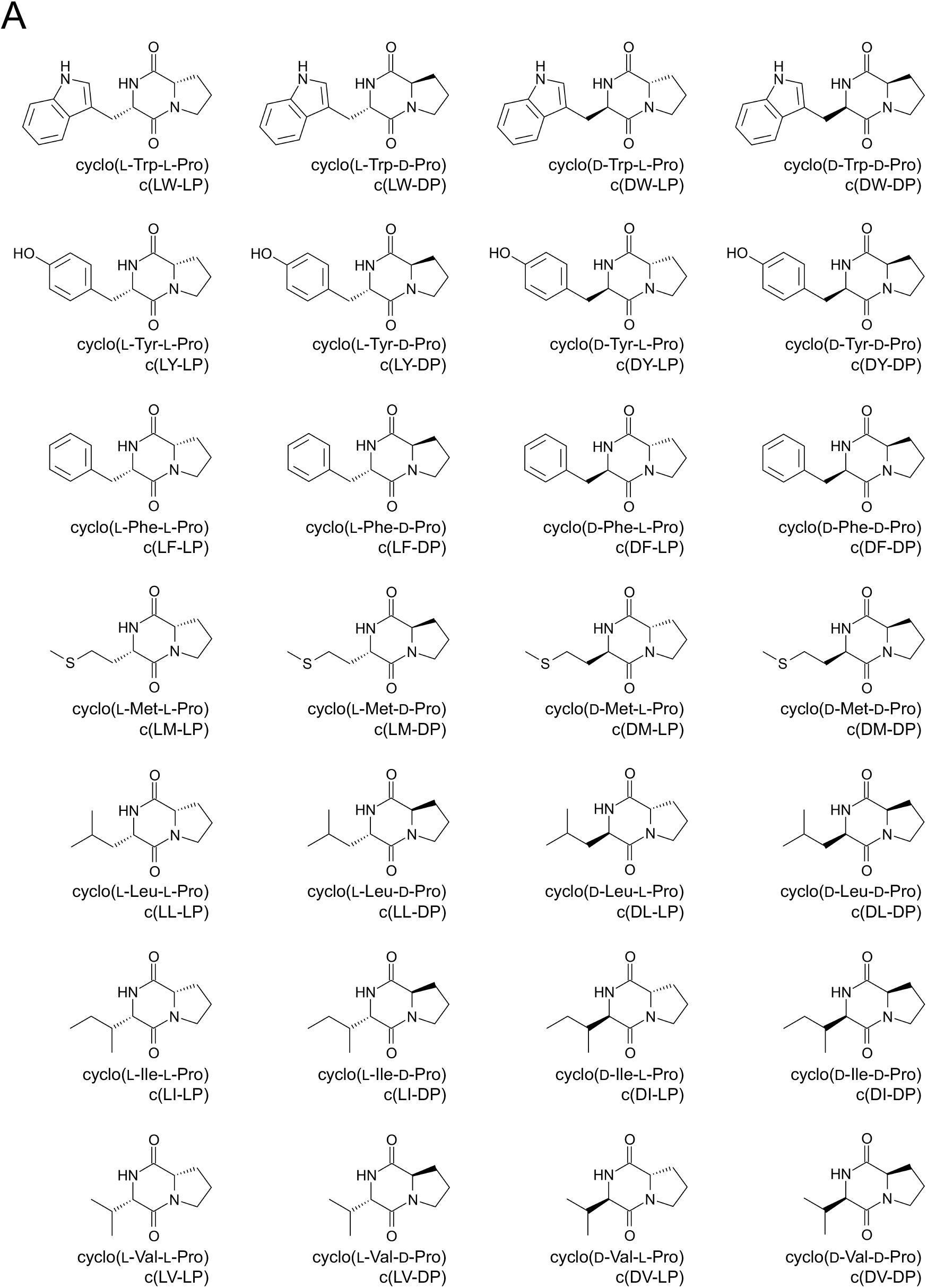
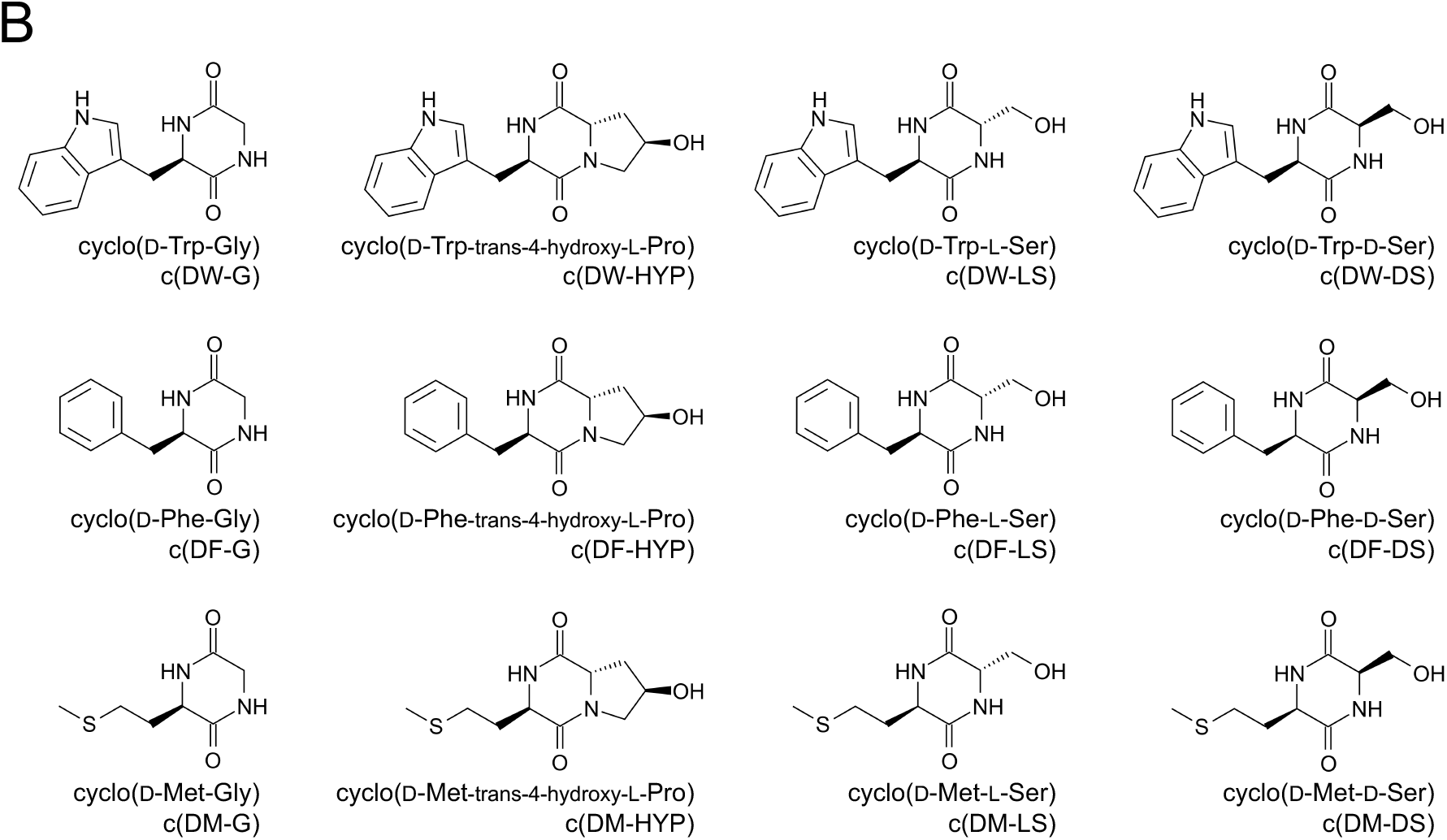
Structures of the chemoenzymatically synthesized DKP library and derivatives. Structures of the 28 DKPs synthesized in a one-pot reaction using the adenylation domain of tyrocidine synthetase A (TycA-A). (B) Structures of derivatives in which the proline residue was replaced with Gly, trans-4-hydroxy-L-Pro, L-Ser, or D-Ser.

**Supplementary Figure S2.**
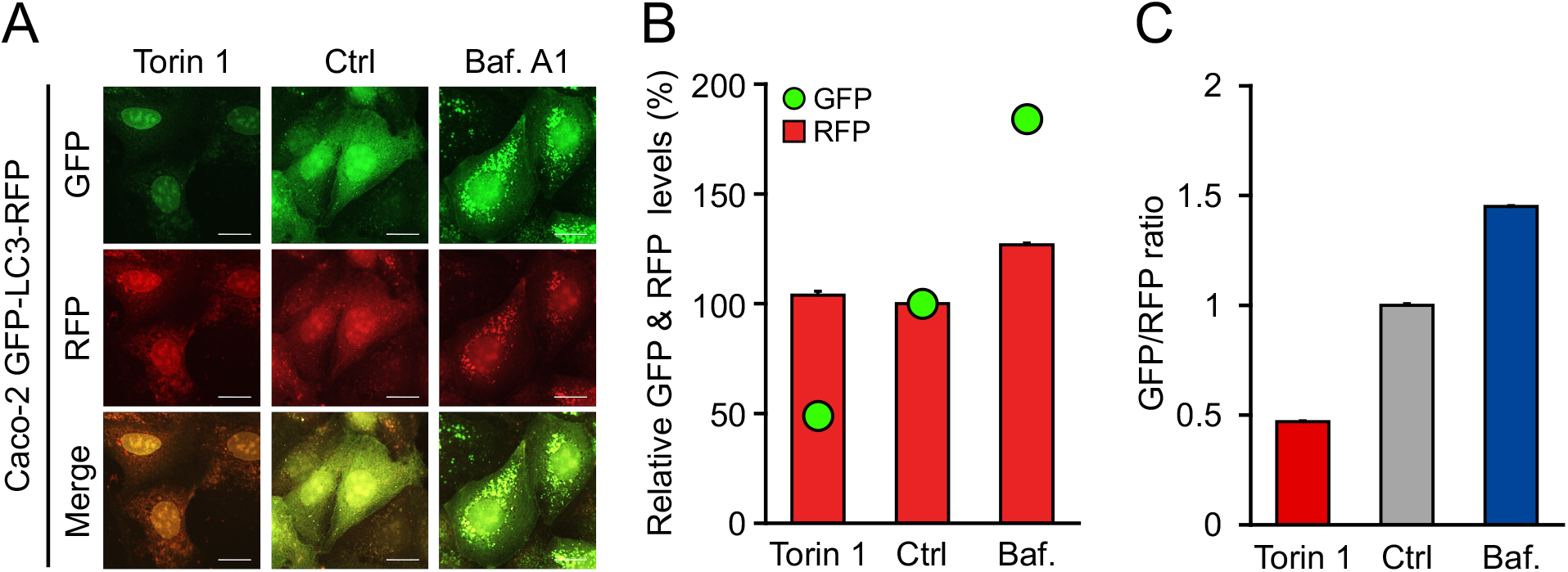
Validation of the GFP-LC3-RFP assay for monitoring autophagic flux. (A) Representative confocal images of Caco-2 cells stably expressing GFP-LC3-RFP after 24 h treatment with 1 μM Torin 1, vehicle control (Ctrl), or 200 nM bafilomycin A1 (Baf. A1). Scale bars, 10 μm. (B) Relative GFP and RFP fluorescence intensities were measured. (C) Quantitative validation by flow cytometry showing decreased GFP/RFP ratios after Torin 1 treatment and increased ratios after bafilomycin A1 (Baf.) treatment. Data are presented as mean ± SEM (n=3).

## References

[1] N. Mizushima, B. Levine, Autophagy in human diseases, N. Engl. J. Med. 383 (2020) 1564–1576, 10.1056/NEJMra2022774.

[2] C. López-Otín, M.A. Blasco, L. Partridge, et al., Hallmarks of aging: An expanding universe, Cell 186 (2023) 243–278, 10.1016/j.cell.2022.11.001.

[3] Y. Aman, T. Schmauck-Medina, M. Hansen, et al., Autophagy in healthy aging and disease, Nat. Aging 1 (2021) 634–650, 10.1038/s43587-021-00098-4.

[4] K.T. Akça, i.Ç. Ayan, S. Çetinkaya, et al., Autophagic mechanisms in longevity intervention: Role of natural active compounds, Expert Rev. Mol. Med. 25 (2023) e13, 10.1017/erm.2023.5.

[5] R.A. Saxton, D.M. Sabatini, mTOR signaling in growth, metabolism, and disease, Cell 168 (2017) 960–976, 10.1016/j.cell.2017.02.004.

[6] A.W. Thomson, H.R. Turnquist, G. Raimondi, Immunoregulatory functions of mTOR inhibition, Nat. Rev. Immunol. 9 (2009) 324–337, 10.1038/nri2546.

[7] D.W. Lamming, L. Ye, P. Katajisto, et al., Rapamycin-induced insulin resistance is mediated by mTORC2 loss and uncoupled from longevity, Science 335 (2012) 1638–1643, 10.1126/science.1215135.

[8] B.L. Kasiske, A. de Mattos, S.M. Flechner, et al., Mammalian target of rapamycin inhibitor dyslipidemia in kidney transplant recipients, Am. J. Transplant. 8 (2008) 1384–1392, 10.1111/j.1600-6143.2008.02272.x.

[9] L. Galluzzi, J.M. Bravo-San Pedro, B. Levine, et al., Pharmacological modulation of autophagy: Therapeutic potential and persisting obstacles, Nat. Rev. Drug Discov. 16 (2017) 487–511, 10.1038/nrd.2017.22.

[10] Q. Zhang, F. Presswalla, R.R. Ali, et al., Pharmacologic activation of autophagy without direct mTOR inhibition as a therapeutic strategy for treating dry macular degeneration, Aging (Albany NY) 13 (2021) 10866–10890, 10.18632/aging.202974.

[11] A.D. Borthwick, N.C. Da Costa, 2,5-Diketopiperazines in food and beverages: Taste and bioactivity, Crit. Rev. Food Sci. Nutr. 57 (2017) 718–742, 10.1080/10408398.2014.911142.

[12] Nagao, Y. Nakamoto, S. Miyauchi, et al., Presence of modified peptides with high bioavailability and angiotensin-converting enzyme inhibitory activity in Japanese fermented soybean paste (Miso), J. Agric. Food Chem. 72 (2024) 18942–18956, 10.1021/acs.jafc.4c02603.

[13] R.M. Huang, X.X. Yi, Y. Zhou, et al., An update on 2,5-diketopiperazines from marine organisms, Mar. Drugs 12 (2014) 6213–6235, 10.3390/md12126213.

[14] A.D. Borthwick, 2,5-Diketopiperazines: Synthesis, reactions, medicinal chemistry, and bioactive natural products, Chem. Rev. 112 (2012) 3641–3716, 10.1021/cr200398y.

[15] K. McCleland, P.J. Milne, F.R. Lucieto, et al., An investigation into the biological activity of the selected histidine-containing diketopiperazines cyclo(His-Phe) and cyclo(His-Tyr), J. Pharm. Pharmacol. 56 (2004) 1143–1153, 10.1211/0022357044139.

[16] F. Fdhila, V. Vázquez, J.L. Sánchez, et al., dd-Diketopiperazines: Antibiotics active against Vibrio anguillarum isolated from marine bacteria associated with cultures of Pecten maximus, J. Nat. Prod. 66 (2003) 1299–1301, 10.1021/np030233e.

[17] S. Nishanth Kumar, C. Dileep, C. Mohandas, et al., Cyclo(D-Tyr-D-Phe): A new antibacterial, anticancer, and antioxidant cyclic dipeptide from Bacillus sp. N strain associated with a rhabditid entomopathogenic nematode, J. Pept. Sci. 20 (2014) 173–185, 10.1002/psc.2594.

[18] L. Wang, Q. Jiang, S. Chen, et al., Natural epidithiodiketopiperazine alkaloids as potential anticancer agents: Recent mechanisms of action, structural modification, and synthetic strategies, Bioorg. Chem. 137 (2023) 106642, 10.1016/j.bioorg.2023.106642.

[19] S. Karakama, S. Suzuki, K. Kino, One-pot synthesis of 2,5-diketopiperazine with high titer and versatility using adenylation enzyme, Appl. Microbiol. Biotechnol. 106 (2022) 4469–4479, 10.1007/s00253-022-12004-y.

[20] T. Kaizuka, H. Morishita, Y. Hama, et al., An autophagic flux probe that releases an internal control, Mol. Cell 64 (2016) 835–849, 10.1016/j.molcel.2016.09.037.

[21] K. Ohnishi, S. Yano, M. Fujimoto, et al., Identification of dietary phytochemicals capable of enhancing the autophagy flux in HeLa and Caco-2 human cell lines, Antioxidants 9 (2020) 1193, 10.3390/antiox9121193.

[22] J. Bojarska, A. Mieczkowski, Z.M. Ziora, et al., Cyclic dipeptides: The biological and structural landscape with special focus on the anti-cancer proline-based scaffold, Biomolecules 11 (2021) 1515, 10.3390/biom11101515.

[23] I.H. Kim, S.-Y. Kim, N.-Y. Park, et al., Cyclo-(L-Phe-L-Pro), a quorum-sensing signal of Vibrio vulnificus, induces expression of hydroperoxidase through a ToxR-LeuO-HU-RpoS signaling pathway to confer resistance against oxidative stress, Infect. Immun. 86 (2018) e00932–17, 10.1128/IAI.00932-17.

[24] H. Jamie, G. Kilian, K. Dyason, et al., The effect of the isomers of cyclo(Trp-Pro) on heart and ion-channel activity, J. Pharm. Pharmacol. 54 (2002) 1659–1665, 10.1211/002235702252.

[25] H. Jamie, G. Kilian, P.J. Milne, Hepatotoxicity of the isomers of cyclo(Trp-Pro), Pharmazie 57 (2002) 638–642, https://pubmed.ncbi.nlm.nih.gov/12369454/.

[26] S.-O. Kang, M.-K. Kwak, Antimicrobial cyclic dipeptides from Japanese quail (Coturnix japonica) eggs supplemented with probiotic Lactobacillus plantarum, J. Microbiol. Biotechnol. 34 (2024) 314–329, 10.4014/jmb.2311.11006.

[27] B.K. Kay, M.P. Williamson, M. Sudol, The importance of being proline: The interaction of proline-rich motifs in signaling proteins with their cognate domains, FASEB J. 14 (2000) 231–241, 10.1096/fasebj.14.2.231.

[28] E. Petrella, L.M. Machesky, D.A. Kaiser, et al., Structural requirements and thermodynamics of the interaction of proline peptides with profilin, Biochemistry 35 (1996) 16535–16543, 10.1021/bi961498d.

[29] R.A. Floto, S. Sarkar, E.O. Perlstein, et al., Small molecule enhancers of rapamycin-induced TOR inhibition promote autophagy, reduce toxicity in Huntington’s disease models and enhance killing of mycobacteria by macrophages, Autophagy 3 (2007) 620–622, 10.4161/auto.4898.

[30] W.K. Martins, M.N. da Silva, K. Pandey, et al., Autophagy-targeted therapy to modulate age-related diseases: Success, pitfalls, and new directions, Curr. Res. Pharmacol. Drug Discov. 2 (2021) 100033, 10.1016/j.crphar.2021.100033.

[31] G.C. Terstappen, C. Schlüpen, R. Raggiaschi, et al., Target deconvolution strategies in drug discovery, Nat. Rev. Drug Discov. 6 (2007) 891–903, 10.1038/nrd2410.

[32] L.P. Nguyen, E.L. Hsu, G. Chowdhury, et al., D-Amino acid oxidase generates agonists of the aryl hydrocarbon receptor from D-tryptophan, Chem. Res. Toxicol. 22 (2009) 1897–1904, 10.1021/tx900043s.

